# Phylogenomic prediction of interaction networks in the presence of gene duplication

**DOI:** 10.1101/2024.08.06.606904

**Authors:** Evan S Forsythe, Tony C Gatts, Linnea E Lane, Chris deRoux, Monica Berggren, Elizabeth A Rehmann, Emily N Zak, Trinity Bartel, Luna L’Argent, Daniel B Sloan

## Abstract

Assigning gene function from genome sequences is a rate-limiting step in molecular biology research. A protein’s position within an interaction network can potentially provide insights into its molecular mechanisms. Phylogenetic analysis of evolutionary rate covariation (ERC) in protein sequence has been shown to be effective for large-scale prediction of functional relationships and interactions. However, gene duplication, gene loss, and other sources of phylogenetic incongruence are barriers for analyzing ERC on a genome-wide basis. Here, we developed *ERCnet*, a bioinformatic program designed to overcome these challenges, facilitating efficient all-vs-all ERC analyses for large protein sequence datasets. We simulated proteome datasets and found that *ERCnet* achieves combined false positive and negative error rates well below 10% and that our novel ‘branch-by-branch’ length measurements outperforms ‘root-to-tip’ approaches in most cases, offering a valuable new strategy for performing ERC. We also compiled a sample set of 35 angiosperm genomes to test the performance of *ERCnet* on empirical data, including its sensitivity to user-defined analysis parameters such as input dataset size and branch-length measurement strategy. We investigated the overlap between *ERCnet* runs with different species samples to understand how species number and composition affect predicted interactions and to identify the protein sets that consistently exhibit ERC across angiosperms. Our systematic exploration of the performance of *ERCnet* provides a roadmap for design of future ERC analyses to predict functional interactions in a wide array of genomic datasets. *ERCnet* code is freely available at https://github.com/EvanForsythe/ERCnet.

## Introduction

Proteins that interact with each other often exhibit correlated rates of sequence evolution across a phylogeny, which have been attributed to shared selective pressures or reciprocal coevolution (Clark et al. 2012). The statistical signature of these interactions is known as evolutionary rate covariation (ERC). ERC analyses have been applied in a wide variety of organisms (Steenwyk et al.; Goh et al. 2000; Ramani and Marcotte 2003; Sato et al. 2005; Clark and Aquadro 2010; De Juan et al. 2013; Wolfe and Clark 2015; Yan et al. 2019; Forsythe et al. 2021; Rei Liao et al. 2022; Tao et al. 2024), and bioinformatic tools have been developed to use ERC to identify novel interaction partners (Wolfe and Clark 2015; Steenwyk et al. 2021) and correlated molecular and phenotypic evolution (Asar et al. 2023; Duchêne et al. 2024). Moreover, ERC analyses are beginning to be applied in a genome-wide manner to yield large sets of predicted protein-protein interactions (i.e, interactome networks) (Steenwyk et al. 2021; Steenwyk et al. 2022). A network view of genetic interactions has proven valuable in revealing the interconnected nature of biological systems (Mao et al. 2009; Rao and Dixon 2019; Wright et al. 2024), and ERC has the potential to be a powerful and efficient method to infer such interaction networks.

As we scale ERC analyses to more taxonomic groups and to larger numbers of genes, phylogenetic incongruence arising from gene duplication/loss, reticulate evolution, and incomplete lineage sorting becomes increasingly problematic (Degnan and Rosenberg 2013; Hahn and Nakhleh 2015). These processes can all lead to trees inferred from different loci (i.e. gene trees) having different branching patterns (i.e., topologies) from each other. Discordant gene trees make it difficult to directly compare evolutionary rates across gene trees because it is unclear which branches correspond to each other. Prior studies have overcome this challenge through combinations of (1) focusing on groups with low levels of phylogenetic incongruence and gene duplication/loss, (2) filtering gene families to retain only gene families with one-to-one orthology, (3) constraining gene tree topologies to the accepted species tree, and (4) measuring branch lengths in a root-to-tip manner that does not depend on topology. These strategies have worked well when the underlying level of phylogenetic incongruence is minimal; however, they may be insufficient in taxonomic groups such as plants that tend to experience more complex genome evolution with extensive gene and whole-genome duplication (Wendel 2015; Panchy et al. 2016; Forsythe et al. 2020). Thus, additional methods are needed to perform ERC analyses in such taxa. A common approach to performing analyses in the face of phylogenetic incongruence is to incorporate gene-tree/species-tree (GT/ST) reconciliation (Vernot et al. 2008; Stolzer et al. 2012; Wu et al. 2014; Comte et al. 2020). GT/ST reconciliation regards phylogenetic incongruence not as noise, but as a signal with which to infer evolutionary events such as gene duplication. Importantly, GT/ST reconciliation also provides a framework for interpreting multiple gene trees in the shared context of a prevailing species tree, which can provide a valuable basis for making direct comparisons between discordant gene trees.

We previously developed a GT/ST reconciliation-based approach to performing genome-wide ERC in plants (Forsythe et al. 2021) that operated at the level of a ‘one-vs-all’ comparison strategy, in which a single locus was tested against a genome-wide set of genes. The biological findings from this approach suggested that large groups of functionally interacting proteins exhibit correlated evolution. However, the limited lens of one-vs-all comparisons only allowed for indirect tests of this hypothesis. A network view of the interactome is needed to assess whether large sets of proteins with correlated rates of evolution share functional relationships. Such a network view requires ERC to be applied in an ‘all-vs-all’ manner that tests all pairwise combinations of proteins. This added dimensionality introduces and exacerbates scaling challenges to ERC analyses.

Here, we present *ERCnet*, a program for performing all-vs-all ERC analyses in the presence of phylogenetic incongruence. We describe our phylogenomic workflow, which automates and parallelizes the computationally intensive analytical steps, and our novel approach to performing branch-by-branch ERC analysis even in trees with gene duplicates. We compile simulated and empirical test datasets to test *ERCnet* performance in response to user-defined analysis parameters and experimental design choices, thus providing an evidence-based roadmap for future ERC analyses.

## Results

### Overcoming gene duplication/loss/incongruence with branch length reconciliation

*ERCnet* (https://github.com/EvanForsythe/ERCnet) is a computational pipeline designed to predict functional interactions among proteins based on phylogenetic signatures of correlated evolution and overcome key scaling challenges associated running all-vs-all analyses on a genome-wide level (Fig. 1). *ERCnet* uses the output of the gene family clustering program *Orthofinder* (Emms and Kelly 2015, 2019), thus making it easy to plug *ERCnet* into existing genomic workflows. Our pipeline employs phylogenetic bootstrap resampling and GT/ST reconciliation strategies to account for phylogenetic incongruence and uncertainty, thus accommodating gene families that have undergone duplication and loss. *ERCnet* automates and parallelizes the computational steps needed to achieve genome-wide network analyses of ERC and consists of four major analytical steps, (1) phylogenomic analyses, (2) GT/ST reconciliation, (3) ERC analyses, and (4) network analyses (See Methods; Fig 1A).

**Fig. 1:**
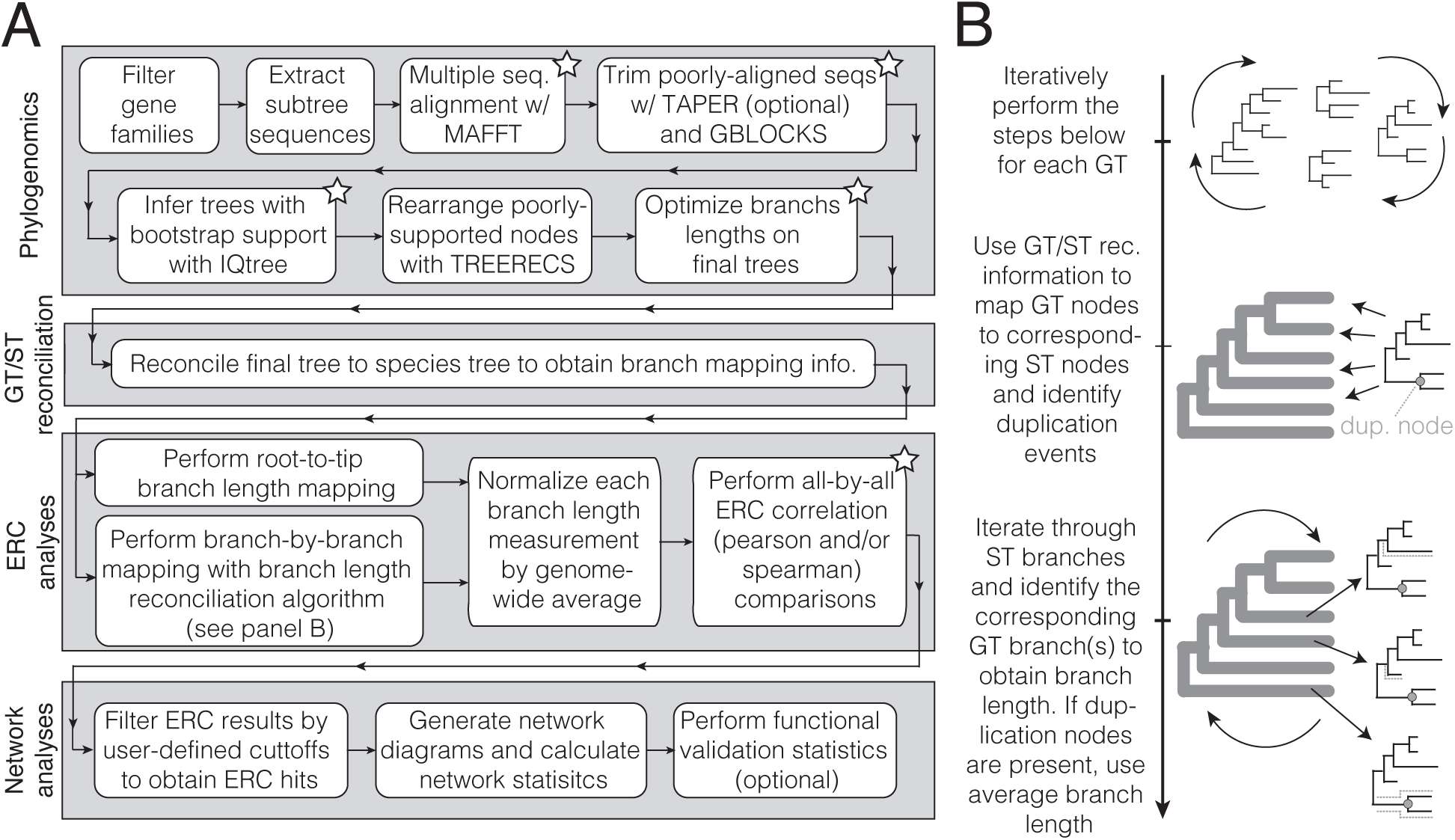
*ERCnet* workflow and algorithm development. (A) Four-step analytical workflow used to analyze *Orthofinder* results (input) and generate an ERC-based interaction network and accompanying summary statistics (output). Stars indicate steps that employ parallel computing. (B) The major analytical steps of the novel ‘branch length reconciliation’ procedure used to calculate branch lengths on a branch-by-branch basis, using species tree (ST) and gene tree (GT) information. Rounded arrows indicate iterative processes.

In addition to automating standard phylogenetic analyses, *ERCnet* provides a novel procedure for comparing rates in a branch-by-branch (BXB) manner even when comparing gene trees with paralogs, missing taxa, and/or topological incongruence. Some previous implementations of ERC have employed a root-to-tip (R2T) methodology for measuring branch lengths. The R2T approach simplifies the task of parsing gene trees (especially when incongruence is present); however, it results in statistical non-independence among datapoints used in correlation analyses because different species can share internal branches within a tree. This statistical shortcoming has been previously recognized (Yan et al. 2019; Forsythe et al. 2021; Smith et al. 2024) but was largely viewed as a ‘necessary evil’ due to lack of a method for traversing discordant gene trees to extract relevant branches for direct comparisons.

Our novel method, branch-length reconciliation (BLR), provides a strategy for avoiding the statistical shortcomings of the R2T method. BLR extracts individual internal and external branch length measurements in a species-tree-aware manner, which allows for direct comparisons across gene trees. The major barrier to performing BXB-based ERC between incongruent phylogenetic trees has been that it is difficult to systematically define which branches (if any) are shared between two gene trees with potentially different taxon-composition and topology. The premise of GT/ST reconciliation methods is that all gene trees evolve within the context of an overarching species tree. This means that the species tree is the common denominator that connects all gene trees in a phylogenomic dataset. BLR uses the species tree as the common comparison point by tallying branch length information from individual gene trees in context of the species tree. BLR accomplishes this task by incorporating the labeled coalescent tree reconciliation structure employed in *DLCpar* (Wu et al. 2014), which provides information that maps gene tree nodes to species tree nodes. Using this information, BLR iteratively extracts the branch(es) on each gene tree that correspond to each species tree branch. During this process BLR also tracks whether a duplication event occurred in the gene tree and, if so, takes the average branch length from the resulting paralogs. When a gene tree lacks information for a given species tree branch, BLR stores an “NA” for that branch. The output of BLR is a table of branch length measurements for each gene tree that are standardized by the species tree branches, allowing comparison across gene trees. This approach provides the ability to extract branch-specific evolutionary rate information from gene trees with incongruent topologies and histories of gene duplication and loss, enabling ERC analyses to include larger gene sets and be applied to a wider range of taxa.

The all-by-all ERC analyses implemented in *ERCnet* yield large tables of ERC results, including p-values calculated from Pearson and/or Spearman correlation analyses. These ERC results are filtered according to user-defined cutoffs to retain the pairwise combinations displaying significant *p*-values and R^2^ values, which we refer to as “ERC hits”. The full set of significant ERC hits form the edges in an interactome network, which can be further analyzed to identify emergent properties, such as clustered modules of cofunctional proteins. Below, we systematically explore the performance of *ERCnet* on simulated and empirical genomic datasets to identify and optimize the parameters that influence computational prediction of interaction networks.

### Accuracy of ERCnet on simulated datasets

We simulated protein sequences for 21 species under a model in which 100 of the 1000 gene families underwent co-accelerated rates of protein evolution (Fig. 2A and B). We used our simulated dataset to assess false positive and negative rates of ERC hits under different *p* and *R^2^* value significance cutoffs and with alternative methodologies for measuring branch lengths (Fig. 2C-F). We found that our BXB method is more conservative than the R2T method and that the Spearman correlation method is more conservative than the Pearson method. The rates obtained from dual filtering (Pearson and Spearman) were very similar to the rates obtained from the Spearman filter alone, meaning Spearman correlation filter acts as the bottleneck in nearly all cases. The combination of BXB and Spearman leads to an especially conservative approach, yielding fewer than 10 total ERC hits and false positive and negative rates approaching 0 and 1 respectively. We summed the false positive and negative rates to identify the overall parameter combination with the lowest combined error rate and found [*BXB; Pearson; P<0.05; R^2^>0.5*] to show the best performance, with a false positive rate of 6.5x10^-4^ (0.065%) and false negative rate of 0. By this metric, the top five error-minimizing combinations all made use of the BXB method and Pearson correlation coefficient, indicating these methods are favorable for minimizing false positive and negative error. The best performing R2T run [*R2T; Pearson; P<0.00001; R^2^>0.2*] yielded a false positive rate of 0.032 (3.2%) and a false negative rate of 0.

**Fig. 2:**
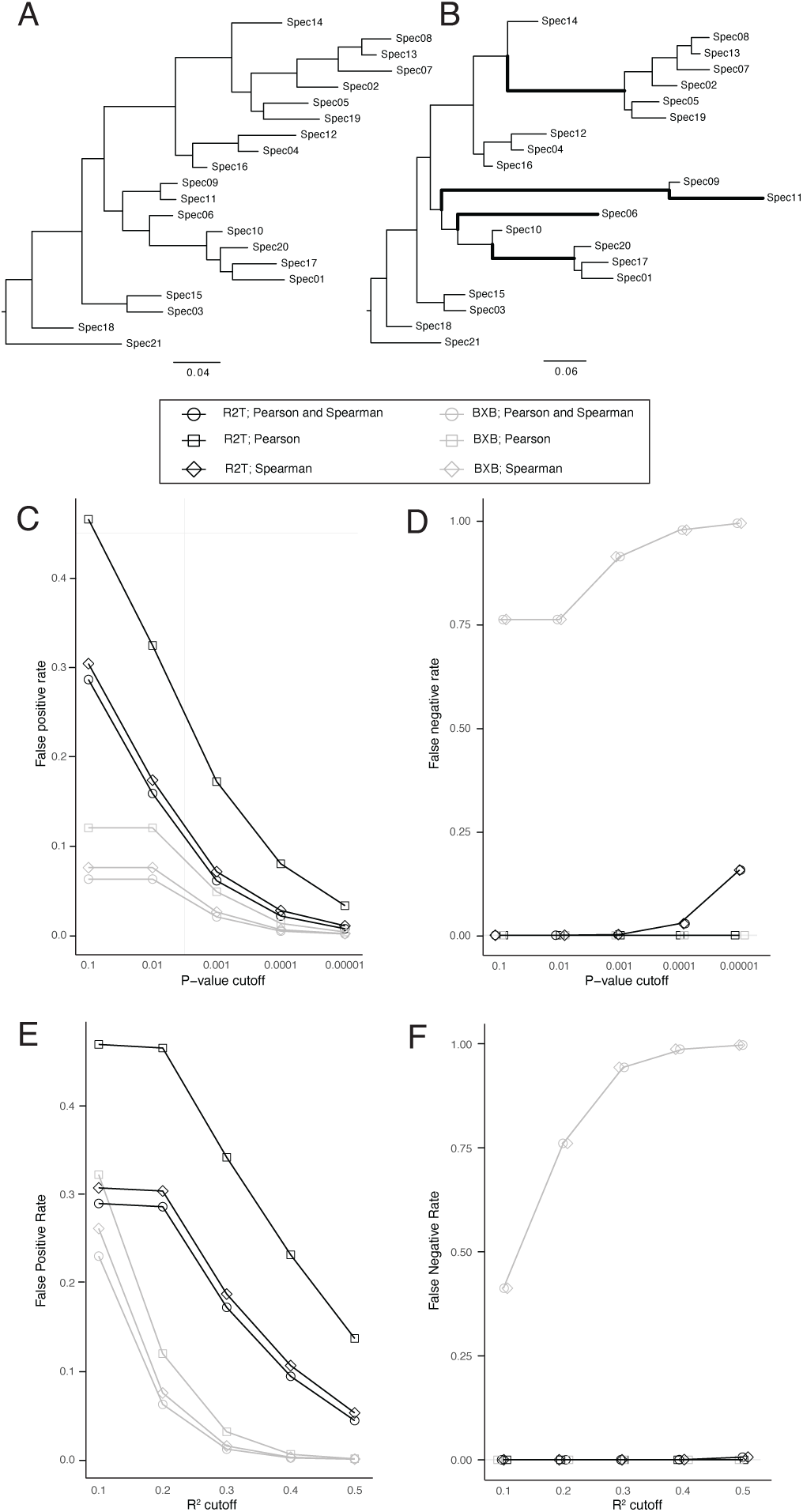
Simulated protein sequences to assess *ERCnet* error rates. (A) The random phylogenetic tree used to simulate the background rates of protein evolution. (B) Tree with the same topology but in which five randomly selected branch lengths (bold branches) were multiplied by 10 in order to simulate co-acceleration for 100 of the 1000 protein families. (C-F) False positive/negative error rates of *ERCnet* runs using different branch length methods (R2T vs BXB) as well as different correlation calculation methods (Pearson vs Spearman) and p-value and R^2^ cutoffs as significance thresholds. Legend shows the branch length method and correlation methods. “Pearson and Spearman” means that ERC hits were only deemed significant if passing filters according to both methods. (C-D) False positive (C) and negative (D) rates assessed across several p-value cutoff values. R^2^ cutoff was held constant at ≥0.20. (E-F) False positive (E) and negative (F) rates assessed across several R^2^ cutoff values. P-value cutoff was held constant at ≤0.05.

Next, we simulated a separate proteome dataset under a model in which rate acceleration takes place in an uncorrelated manner, meaning any ERC hit obtained from this dataset would be considered an ‘spurious’ ERC hit (Fig. S1). Consistent with the results above, BXB performed in a far more conservative manner compared to R2T on the non-accelerated dataset. R2T showed more than 60,000 spurious hits, whereas the highest number of spurious hits in a BXB run was 19,395. Taken together, these results show that *ERCnet* achieves very promising levels of accuracy on simulated data and that our BXB method will be a valuable tool in minimizing error in ERC analyses.

### Proteome coverage and statistical power for ERCnet analyses of sample datasets

ERC has been applied in a variety of organisms with considerable success. However, a systematic evidence-based exploration of the power, accuracy, and efficiency of the ERC method on real genomic datasets has never been performed. A consequential decision in ERC analyses is the number of species to include. Currently, researchers have little guidance in making taxon-sampling choices for ERC analyses, resulting in somewhat arbitrary choices that can likely have a substantial impact on ERC performance. To address this limitation, we compiled a set of 35 plant proteomes (i.e., protein sequences from all annotated genes in the genome), spanning angiosperm diversity (Table S1). We randomly subsampled sets of these proteomes to create test datasets of varying sizes. The resulting datasets contained *n*=10, *n*=15, *n*=20, and *n*=25 in-group taxa (see Methods). Five random replicates were performed for each *n* value, resulting in a total of 20 datasets, each of which was subjected to full *ERCnet* analyses (Fig. 3).

**Fig. 3:**
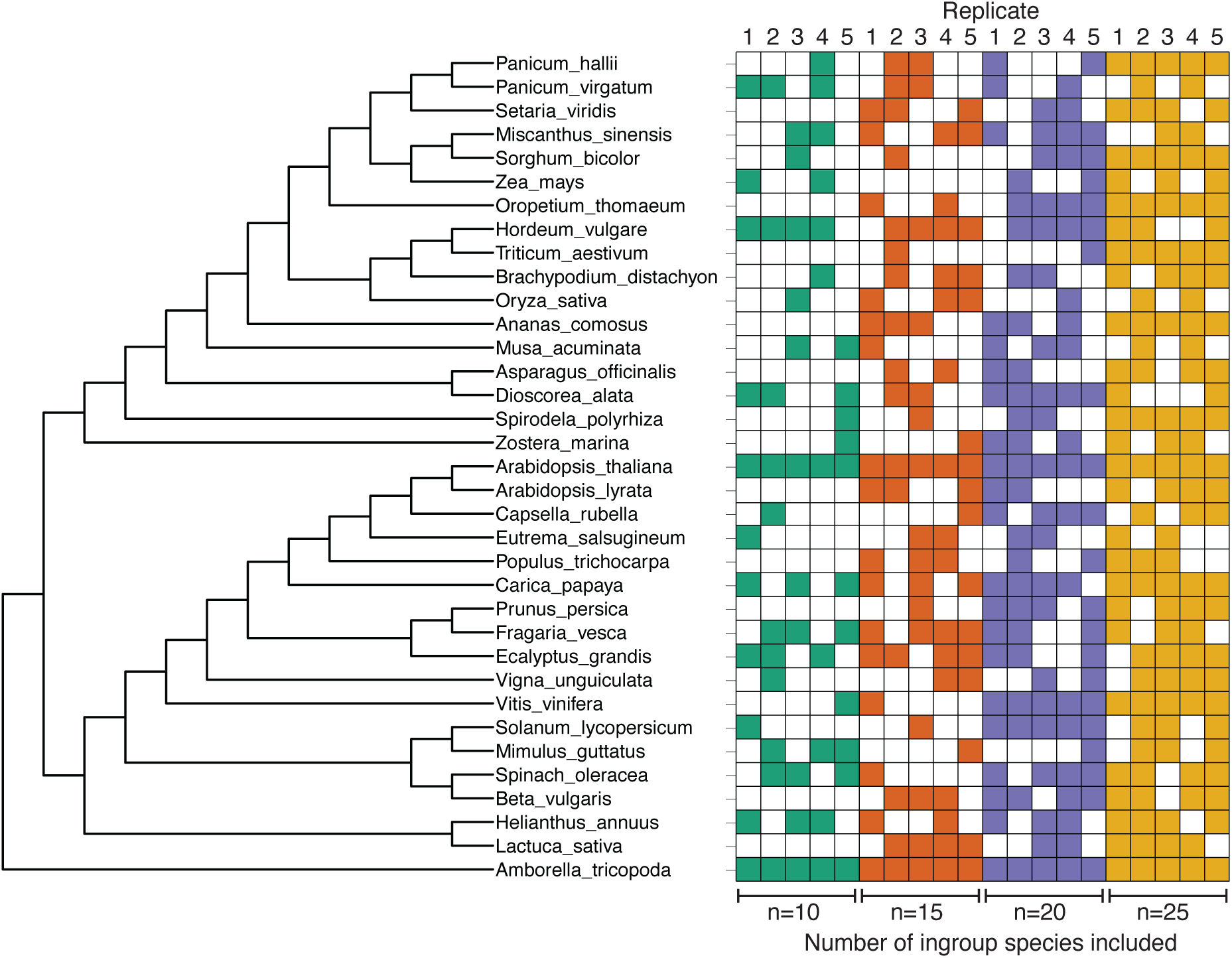
Angiosperm taxon-sampling datasets used to assess *ERCnet* performance. (Left) Phylogenetic tree of the full pool of species included for random sub-sampling. (Right) Presence-absence plot indicating the species included in random datasets of each size. Five replicates were performed for each dataset size. *Amborella trichopoda* was included as an outgroup for all replicates and *Arabidopsis thaliana* was included as a common ingroup representative for all replicates.

The first step of *ERCnet* is to filter *Orthofinder* data to identify the gene families (“orthogroups”) that are suitable for phylogenomic analyses by applying filtering cutoffs based on minimum number of species represented and maximum number of gene copies per species. *ERCnet* provides a parameter scan option for users to make an informed choice on filtering settings (see Methods). To understand how filtering parameters affect dataset size, we applied a filtering formula across all datasets (see Methods) and found that datasets with a larger number of species yield a smaller number of genes retained after filtering (Fig. 4A). While datasets with a larger number of species tend to have more total gene families, these gene families tend to be filtered out at a disproportionally high rate based on both the minimum species representation filter and the maximum number of gene copies per species filter (Table S2), leading to a net loss of proteome coverage at larger dataset sizes.

**Fig. 4:**
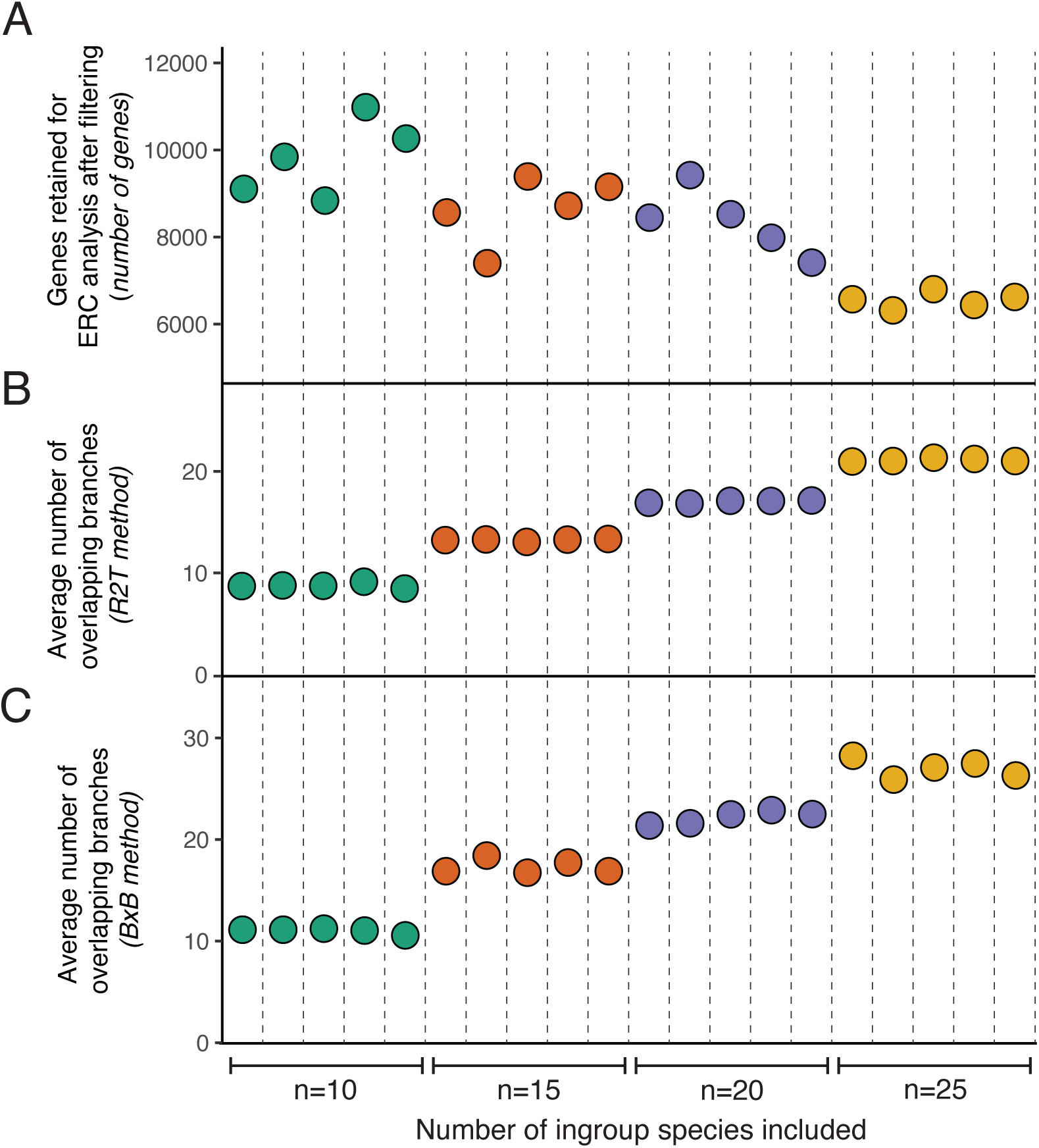
Proteome coverage and overlapping branches after *ERCnet* filtering. (A) The number of proteins retained for phylogenomic analysis after several phases of quality-control filtering that occur during the first steps of *ERCnet*. These number represents the number of proteins that are tested for interaction during ‘all-vs-all’ ERC analyses at later steps of *ERCnet.* (B-C) The number of overlapping branches (i.e. points on correlation plots) among pairs of proteins during all-vs-all ERC analyses. For the root-to-tip method (B), ‘branches’ refers to paths from root of tree to each tip. For the branch-by-branch method (C), ‘branches’ refers to the common branches determined by our ‘branch length reconciliation’ method.

The gene families that pass initial filtering are next run through the phylogenomic pipeline and ultimately subjected to all-vs-all ERC correlation analyses. Each pairwise ERC comparison between two gene trees constitutes a correlation analysis. The number of points for each correlation corresponds to the number of branches shared between the two gene trees being compared, which is variable because each gene may have experienced different degrees of gene duplication and loss. The number of overlapping branches is also dependent on the method used to measure branch lengths (*i.e.* R2T vs BXB). We found that datasets with larger number of species yield a higher average number of overlapping branches (Fig. 4B and C). This result is expected because larger datasets yield larger gene trees, which inherently have more branches. Comparing R2T results (Fig. 4B) to BXB results (Fig. 4C), we see that BXB tends to yield a higher number of overlapping branches, which stems from including internal branches under BXB, while the R2T methodology is limited to the number of species in the in-group. For example, a gene tree with four ingroup species will only have four R2T measurements but will have six branches when internal branches are included. In general, the number of overlapping branches will directly impact the sample size of correlation analyses, suggesting that BXB has the potential to add statistical power to ERC analyses. Thus, there is a clear tradeoff between the proportion of the genome that can be probed with *ERCnet* (Fig. 4A) and the potential statistical power of ERC analyses (Fig. 4B and C).

### Size and functional clustering of networks across datasets

We used our simulation results (Fig. 2 and S1) to guide our choice of significance filtering criteria to use on our empirical datasets. We chose [*Pearson; P<0.0001; R^2^>0.4*] to obtain a reasonable balance between false positive and negative error across both R2T and BXB-based networks. We applied this filter to obtain an interactome network for each *ERCnet* run (Fig. 5, S2, and S3). In general, we found that R2T analyses returned a much higher number of ERC hits than BXB analyses, resulting in larger networks in terms of both nodes and edges (Fig. 5A vs B). The large number of hits observed for R2T analyses could partially result from pseudoreplication due to statistical non-independence of internal branch lengths (described above). Another general pattern is that networks show more variability between replicates for lower *n*-values. This is likely at least partially driven by randomness playing a larger role at smaller sample sizes in our taxon sampling from a fixed-size pool (Fig. 3), highlighting that selection of individual species has a large impact on ERC-based networks at small *n*-values. Interestingly, we do not detect clear trends between *n* and the size of networks. For R2T-based networks, there is some evidence that small *n*-values yield networks with fewer edges (Fig. 5A), consistent with idea that R2T-based methods may not achieve a sufficient number of overlapping branches to yield significant ERC hits (see Fig. 4B). BXB ERC analyses, on the other hand, display the highest number of ERC hits for intermediate *n* values (Fig 5B). This result suggests that there could be a ‘sweet-spot’ that balances the number of genes retained after filtering (see Fig. 4A) and the number of overlapping branches (see Fig. 4C) to achieve coverage and statistical power for BXB-based ERC analyses. Taken together, the differences in trends for R2T versus BXB-based analyses indicate that the optimal dataset size depends on the branch-length measuring method.

**Fig. 5:**
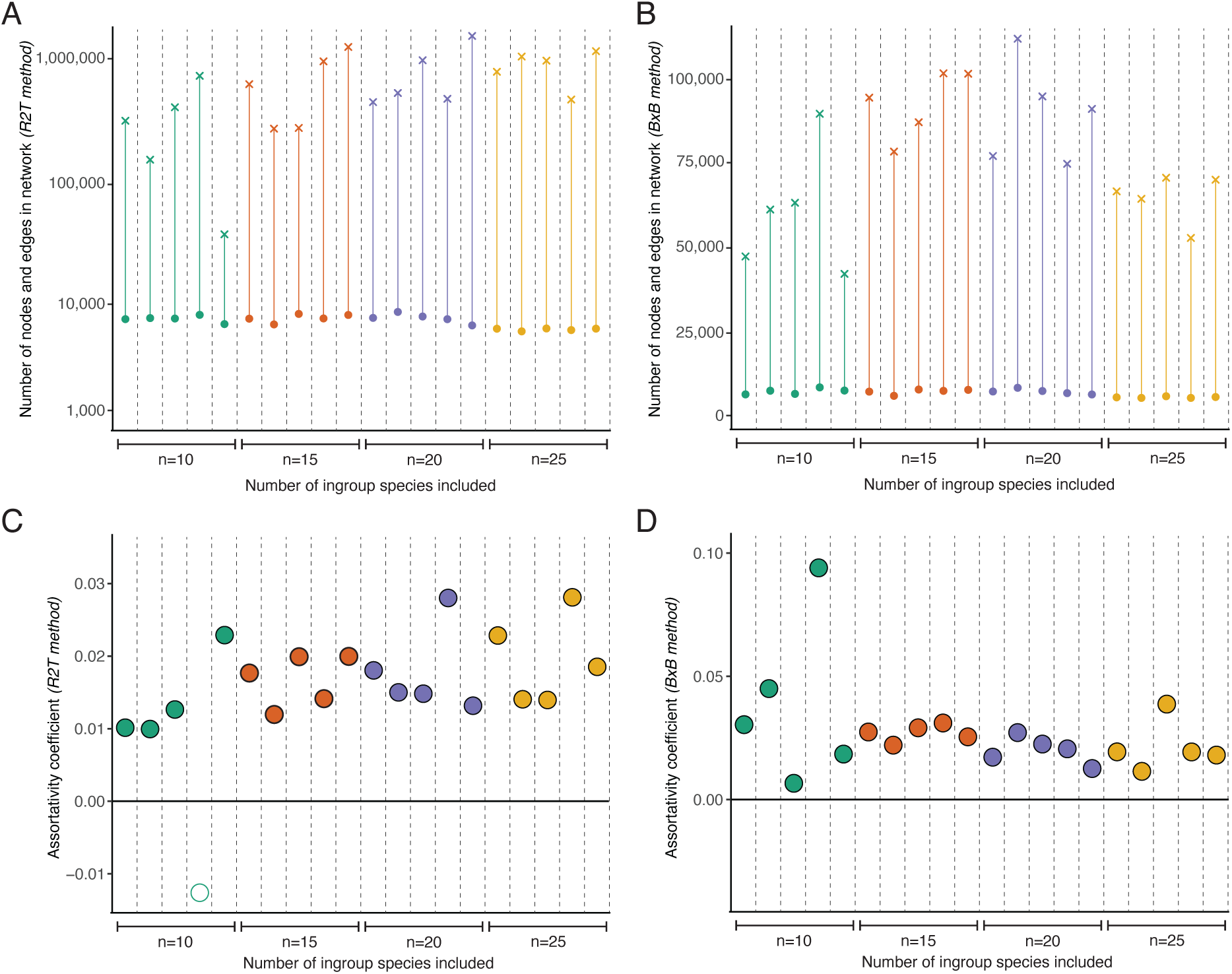
Composition and functional clustering of *ERCnet* interaction networks. (A-B) The number of nodes (points) and edges (x’s) in networks obtained using the root-to-tip (A) and branch-by-branch method (B). Note the log scale for panel A. (C-D) The assortativity coefficient estimated from. Filled points indicate the assortativity coefficient is significantly greater than the randomized null distribution (z-score from randomized permutation test). Significant positive assortativity indicates clustering of traits across a network. The trait measured here was the predicted targeting (plastid, mitochondrial, other) of the proteins in the network.

The number of ERC hits yielded by *ERCnet* analyses defines the total number of edges in the interaction network output with each run of *ERCnet*. The size of the network is an important component of *ERCnet* performance; however, it is also important to understand contribution of biological signal versus noise in empirical ERC networks. We sought a summary statistic that can serve as a proxy for biological signal in ERC networks. To this end, we incorporated subcellular localization annotations from *Arabidopsis thaliana* and used the graph-theory assortativity coefficient to quantify the degree to which co-localized proteins are clustered in interaction networks (Fig. 5C-D). For BXB analyses, we found that all but one of the *ERCnet* runs yielded networks with significantly positive assortativity coefficient, providing another promising indication that *ERCnet* successfully detects ‘true’ biological signal at a level that outweighs background noise. BXB shows significantly positive assortativity across all replicates (Fig 5D). There is not a clear trend between *n* and clustering, but, similar to network size (Fig 5A-B), there is much more variability between replicates at *n*=10. R2T shows a slight positive relationship between *n* and assortativity. This includes a clear outlier replicate [n=10; replicate=4] that shows negative assortativity coefficient, suggesting that noise and error outweigh biological signal under this combination of species and the R2T branch length method. These results highlight that internal biological validation will play an important role in interpreting ERC-based interaction networks.

### Overlap in interactions predicted across multiple datasets

In addition to understanding how different datasets and parameters affect the number of ERC hits produced by *ERCnet*, it is also important to understand the consistency with which individual protein-protein interactions are predicted across different datasets. Therefore, we compared the overlap of specific *ERCnet* hits (i.e. pairs of proteins) across different taxon sampling datasets (Fig. 6). We found that there were no ERC hits that persisted across all 20 of the R2T and BXB *ERCnet* runs. More than half of the ERC hits runs were non-overlapping ‘singletons’ (i.e. found in a single *ERCnet* run); however, there were hits with a degree overlap was as high as 19-way overlap and 15-way overlap among R2T and BXB *ERCnet* runs, respectively. We might expect a background level of overlap, even if ERC hits were driven entirely by noise, so we generated randomized replicates in which the identities of the interaction partners were randomly swapped within each *ERCnet* run (Fig. 6, grey bars). We found that the observed level of overlap exceeds a randomized background, with far more ERC hits occupying rightward tail of the distribution (high degree of overlap) in our observed data. This result shows that ERC signal rises above background noise and persists across *ERCnet* runs that use different input data. However, the general orientation of distributions toward low-degree overlap highlights the importance of species sampling on *ERCnet* results.

**Fig. 6:**
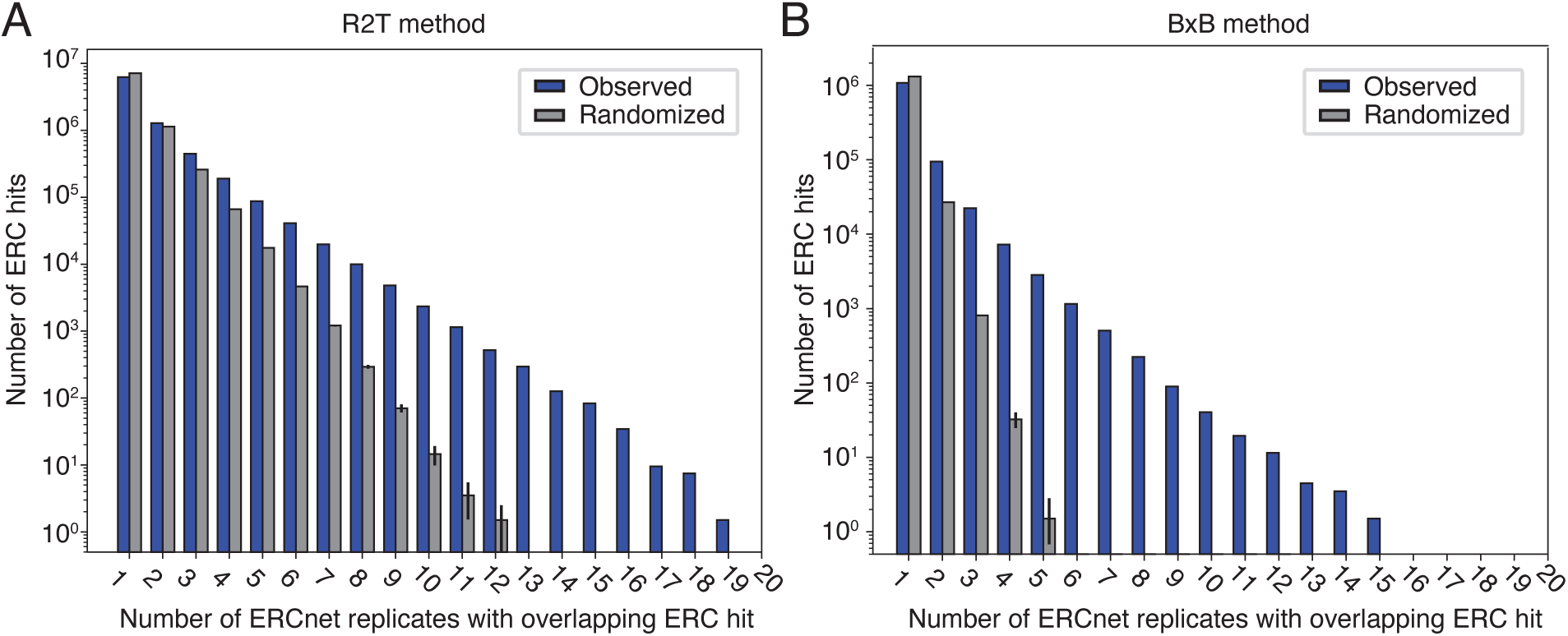
Overlap in *ERCnet* hits between runs. (A-B) Bar plots describing the number of hits plotted by the number of *ERCnet* replicate runs they appeared in. Blue bars show observed values, and gray bars show averages from ten randomized replicates. Lines indicate standard error from randomized replicates.

Moreover, our sampling approach provides a unique opportunity to identify pairs of proteins that show a consensus signature of correlated evolution across *ERCnet* runs. We compiled the ERC hits from the three highest levels of overlap for R2T and BXB runs (Table 1). This shortlist of ‘consensus’ ERC hits contains 17 R2T consensus hits and eight BXB consensus hits. Using the *Arabidopsis thaliana* homolog from each gene family, we obtained gene names, subcellular localization information, and descriptions and from *The Arabidopsis Information Resource* (TAIR) (Lamesch et al. 2012). We found that 8/17 and 5/8 ERC hits show evidence of subcellular co-localization for R2T and BXB runs, respectively. 9/17 and 2/8 ERC hits show at least some evidence of co-functionality based on gene descriptions for R2T and BXB runs, respectively. Taking co-localization and co-functionality information together, we find that 10/17 and 6/8 ERC hits show evidence of plausible interaction by at least one of these annotation metrics for R2T and BXB runs, respectively.

**Table 1:** Consensus ERC hits from angiosperm taxon-sampling datasets. ERC hits found in the top three categories of overlap for BXB or R2T *ERCnet* runs. (**Meth/Overlap.**) The branch length method and a number of overlapping *ERCnet* runs the hit was found in parentheses. (**AGI[s]**) Presence of Multiple Arabidopsis Gene Identifiers (AGIs) indicates that multiple Arabidopsis paralogs were present in the gene family. In cases of multiple paralogs, we used to first one (underlined) to obtain TAIR information. (**Name and Description**) Gene name and description from TAIR. (**Loc.)** The subcellular localization as indicated by ‘Cellular Component’ Gene Ontology terms. (**Loc. evid.**) Indicates whether the “Loc.” columns show at least some overlap between Gene A and Gene B *(Yes/No/NA)*. (**Desc. evid.**) Qualitative assessment of whether the information in ‘Description’ columns points to cofunctionality/interaction between Gene A and Gene B *(None, Weak, Strong, Unknown)*. ‘Unknown’ indicates lack of sufficient information in at least one of the two genes.

| Meth./<br>Overlap. | Gene A |  |  |  | Gene B |  |  |  | Loc. evid. | Desc. evid. |
| --- | --- | --- | --- | --- | --- | --- | --- | --- | --- | --- |
|  | AGI(s) | Name | Description | Loc. | AGI(s) | Name | Description | Loc. |  |  |
| R2T<br>(19) | AT1G09130 | CLPR3 | ATP-dependent caseinolytic (Clp) protease/crotonase family protein | chloro., mito. | AT2G27460 | ATSEC23D | Sec23 homolog, forms a distinct clade with SEC23A. Mutants have defects in pollen exine patterning, tapetal development and pollen intine formation | cytoplasm, endoplasmic reticulum exit site | No | None |
| R2T<br>(18) | AT1G09130 | CLPR3 | ATP-dependent caseinolytic (Clp) protease/crotonase family protein | chloro., mito. | AT3G44890 | RIBOSOMAL PROTEIN BL9C | Plastid ribosomal protein CL9. The mRNA is cell-to-cell mobile | chloro., cytos. | Yes | Strong |
| R2T<br>(18) | AT1G29340 | PLANT U-BOX 17 | Encodes a protein containing a UND, a U-box, and an ARM domain. This protein has E3 ubiquitin ligase activity. It is required for cell death and full resistance specified by Arabidopsis RPM1 and RPS4 resistance proteins against *Pseudomonas syringae* pv tomato. The mRNA is cell-to-cell mobile | chloro. | AT1G68020 | TREHALOSE-6-PHOSPHATASE SYNTHASE S6 | Encodes an enzyme putatively involved in trehalose biosynthesis. The protein has a trehalose synthase (TPS)-like domain and a trehalose phosphatase (TPP)-like domain. It can complement a yeast mutant lacking both of these activities suggesting that this is a bifunctional enzyme | chloro. | Yes | None |
| R2T<br>(18) | AT1G14000 | VH1-INTERACTING KINASE | Encodes a protein with similarity to members of the C1 subgroup of MAP kinase kinase kinases. Interacts physically with the receptor kinase BRL2/VH1 and appears to be involved in auxin and brassinosteroid signaling. The mRNA is cell-to-cell mobile. | extracellular, nuc., vacuole | AT2G27460 | ATSEC23D | Sec23 homolog, forms a distinct clade with SEC23A. Mutants have defects in pollen exine patterning, tapetal development and pollen intine formation | cytoplasm, ER exit site | No | Weak |
| R2T<br>(18) | AT1G06440 | NA | Ubiquitin carboxyl-terminal hydrolase family protein | chloro. | AT5G63100 | NA | S-adenosyl-L-methionine-dependent methyltransferases superfamily protein | chloro., mito. | Yes | Weak |
| R2T<br>(18) | AT3G49730<br>AT5G65820 | NA | Tetratricopeptide repeat (TPR)-like superfamily protein | mito. | AT4G14590 | DEFECTIVE IN SNRNA PROCESSING 2, EMBRYO DEFECTIVE 2739 | embryo defective 2739 | nuc. | No | None |
| R2T<br>(18) | AT1G11750 | CLPP6 | One of several nuclear-encoded ClpPs (caseinolytic protease). Contains a highly conserved catalytic triad of Ser-type proteases (Ser-His-Asp) | chloro. | AT3G44890 | RIBOSOMAL PROTEIN BL9C | Plastid ribosomal protein CL9. The mRNA is cell-to-cell mobile. | chloro., cytos. | Yes | Strong |
| R2T<br>(18) | AT1G09130 | CLPR3 | ATP-dependent caseinolytic (Clp) protease/crotonase family protein | chloro., mito. | AT1G49970 | CLPR1 | Encodes a ClpP-related sequence. Though similar to ClpP proteins, this does not contain the highly conserved catalytic triad of Ser-type proteases (Ser-His-Asp) | chloro., nuc. | Yes | Strong |
| R2T<br>(17) | AT1G49970 | CLPR1 | Encodes a ClpP-related sequence. Though similar to ClpP proteins, this does not contain the highly conserved catalytic triad of Ser-type proteases (Ser-His-Asp) | chloro., nuc. | AT5G09995 | REGULATOR OF FATTY ACID SYNTHESIS 1 | Regulates de novo fatty acid synthesis by modulating hetACCase distribution. | chloro. | Yes | Weak |
| R2T<br>(17) | AT1G09130 | CLPR3 | ATP-dependent caseinolytic (Clp) protease/crotonase family protein | chloro., mito. | AT1G11750 | CLPP6 | One of several nuclear-encoded ClpPs (caseinolytic protease). Contains a highly conserved catalytic triad of Ser-type proteases (Ser-His-Asp) | chloro. | Yes | Strong |
| R2T<br>(17) | AT1G29340 | PLANT U-BOX 17 | Encodes a protein containing a UND, a U-box, and an ARM domain. This protein has E3 ubiquitin ligase activity. It is required for cell death and full resistance specified by Arabidopsis RPM1 and RPS4 resistance proteins against *Pseudomonas syringae* pv tomato. The mRNA is cell-to-cell mobile | chloro. | AT2G27460 | ATSEC23D | Sec23 homolog, forms a distinct clade with SEC23A. Mutants have defects in pollen exine patterning, tapetal development and pollen intine formation | cytoplasm, ER exit site | No | None |
| R2T<br>(17) | AT4G14590 | DEFECTIVE IN SNRNA PROCESSING 2 | embryo defective 2739 | Nuc. | AT5G50280 | ECD2, EMB1006 | Responsible for chloroplast gene expression and group II intron splicing of several genes. Associated with the expression of ribosomal genes and accumulation of chloroplast ribosomes. Critically important for early chloroplast development in cotyledon | chloro. | No | Strong |
| R2T (17) | AT2G27460 | ATSEC23 D | Sec23 homolog, forms a distinct clade with SEC23A. Mutants have defects in pollen exine patterning, tapetal development and pollen intine formation. | cytop., ER exit site | AT3G49730<br>AT5G65820 | NA | Tetratricopeptide repeat (TPR)-like superfamily protein; (source: Araport11) | mito. | No | None |
| R2T (17) | AT1G14000 | VH1-INTERACTING KINASE | Encodes a protein with similarity to members of the C1 subgroup of MAP kinase kinase kinases. Interacts physically with the receptor kinase BRL2/VH1 and appears to be involved in auxin and brassinosteroid signaling. The mRNA is cell-to-cell mobile | extracell., nuc., vacuole | AT4G29060 | EMBRYO DEFECTIVE 2726 | Involved in chloroplast biogenesis and early embryo development. May function as an EF-Ts to regulate plastid translation | chloro., plasma memb. | No | None |
| R2T (17) | AT1G33340 | PICALM 8 | ENTH/ANTH/VHS superfamily protein | clathrin-coated vesicle | AT1G68020 | TREHALOSE-6-PHOSPHATASE SYNTHASE 6 | Encodes an enzyme putatively involved in trehalose biosynthesis. The protein has a trehalose synthase (TPS)-like domain and a trehalose phosphatase (TPP)-like domain. It can complement a yeast mutant lacking both of these activities suggesting that this is a bifunctional enzyme | Chloro. | No | None |
| R2T (17) | AT1G09130 | CLPR3 | ATP-dependent caseinolytic (Clp) protease/crotonase family protein | chloro., mito. | AT4G17560<br>AT5G47190 | RIBOSOMAL PROTEIN BL19CZ | Ribosomal protein L19 family protein | chloro., cytos. | Yes | Strong |
| R2T (17) | AT1G09130 | CLPR3 | ATP-dependent caseinolytic (Clp) protease/crotonase family protein | chloro., mito. | AT3G58800 | NA | secretion-regulating guanine nucleotide exchange factor | nuc. | No | None |
| BXB (15) | AT1G29850 | PROGRAMMED CELL DEATH PROTEIN 5 | Encodes a protein that by its interaction with HAM acetyltransferases plays an important role during DNA damage responses induced by UV-B radiation and participates in programmed cell death programs | cytos., nuc. | AT5G60340 | ADENYLATE KINASE 6 | Encodes a nuclear adenylate kinase that interacts with a putative homolog of Rps14, AtRPS14-1 and affects the elongation of cells in the stem | nuc. | Yes | None |
| BXB (14) | AT4G36580<br>AT2G18330 | SHOT1 BINDING ATPASE 4 | Homologue of animal ATPase Family AAA Domain-Containing Protein 3 (ATAD3), which is involved in mitochondrial nucleoid organization; interacts with SHOT1 | plasma memb. | AT5G16930<br>AT3G03060 | SHOT1 BINDING ATPASE 2 | Homologue of animal ATPase Family AAA Domain-Containing Protein 3 (ATAD3), which is involved in mitochondrial nucleoid organization; interacts with SHOT1 | chloro., mito. | No | Strong |
| BXB (14) | AT1G51110 | FIBRILLIN10 | PAP/fibrillin (ECM1) localized to chloroplasts; involved in structural activity | chloro., nuc. | AT5G27560 | DUF1995 domain protein | NA | chloro. | Yes | Unknown |
| BXB (14) | AT3G09150 | GENOMES UNCOUPLED 3 | Required for biosynthesis of the tetrapyrrole phytochrome chromophore phytochromobilin. Encodes phytochromobilin synthase, a ferredoxin-dependent biliverdin reductase. It is necessary for coupling the expression of some nuclear genes to the functional state of the chloroplast | chloro. | AT5G27560 | DUF1995 domain protein | NA | chloro. | Yes | Unknown |
| BXB (13) | AT1G51110 | FIBRILLIN10 | PAP/fibrillin (ECM1) localized to chloroplasts; involved in structural activity | chloro., nuc. | AT5G64670 | RIBOSOMAL PROTEIN UL15M | Ribosomal protein L18e/L15 superfamily protein | mito. | No | None |
| BXB (13) | AT1G05385 | LOW PSII ACCUMULATION 19 | Encodes a Psb27 homolog involved in photosystem II biogenesis | chloro., cyto. | AT1G51110 | FIBRILLIN10 | PAP/fibrillin (ECM1) localized to chloroplasts; involved in structural activity | chloro., nuc. | Yes | Weak |
| BXB (13) | AT1G51110 | FIBRILLIN10 | PAP/fibrillin (ECM1) localized to chloroplasts; involved in structural activity | chloro., nuc. | AT4G27700 | NA | Rhodanese/Cell cycle control phosphatase superfamily protein | chloro., cytoskel. | Yes | None |
| BXB (13) | AT1G10790 | NA | hydroxyproline-rich glycoprotein family protein | NA | AT5G25640 | NA | Rhomboid-related intramembrane serine protease family protein | NA | NA | Unknown |

A dominant pattern from both R2T and BXB consensus ERC hits is that chloroplast-localized proteins appear to be especially prevalent. 12/13 ERC hits that exhibit co-localization share localization in the chloroplast, suggesting that there are fundamental dynamics to chloroplast biology that drives consistent patterns of ERC across a variety of angiosperm datasets. Among the R2T consensus hits, members of the plastid caseinolytic protease (Clp) complex are especially prevalent; we observe two ERC hits among members of the Clp complex *sensu stricto* as well as three ERC hits between a Clp protein and a plastid ribosomal protein. These results are consistent with prior results, in which R2T and related methods revealed strong ERC among Clp and ribosomal proteins (Forsythe et al. 2021; Gatts et al. 2024). However, it is notable that previous studies specifically sampled species known *a priori* to exhibit accelerated plastid genome evolution, whereas the species sampled here chosen without regard to plastid evolutionary rates. Therefore, the results shown here provide the new insight that ERC signatures spanning the plastid proteostasis apparatus are prevalent in a more general and widespread sense in angiosperms. Similar to R2T consensus hits, BXB consensus hits appear to be enriched for organelle-localized proteins. However, they otherwise lack a clear pattern of functional enrichment, instead touching on several areas of plant cellular biology such as cellular signaling, the cell cycle, chloroplast and mitochondrial organization, and retrograde signaling.

### Parallel processing and computational runtime

Scaling ERC analyses to whole proteomes involves computationally intensive statistical steps that are best suited for high-performance computing (HPC). *ERCnet* provided built-in and customizable parallelization at the most computationally intensive steps, allowing users to tailor *ERCnet* to perform optimally in a variety of computing environments. We ran *ERCnet* on our test datasets on a server, utilizing 48 parallel threads (Fig 7). Through our early testing of *ERCnet*, we found that the pairwise ERC analysis steps (implemented with ERC_analyses.py) suffered a performance deficit when run on more than 24 parallel threads, perhaps due to a parallel computing phenomenon known as ‘thrashing’, in which increased resource allocation comes at an overhead cost for the operating system to schedule, resulting in reduced efficiency. Therefore, we run this step using only 24 threads. Total runtimes for full *ERCnet* analyses ranged from ∼12-43 hours. We find that all-by-all ERC step constitutes a larger portion of the total runtime step at *n*=10, whereas the phylogenomic analyses step rules the runtime at larger *n-*values (Fig. 7A). Somewhat counterintuitively, *n*=10 runs exhibited some of the longest total runtimes. These patterns are likely a product of the complex relationship between proteome coverage after filters, size of individual genes trees, and the fact that processing in the phylogenomics step scale linearly with number of genes in the dataset, while all-by-all ERC analyses is proportional to the square of the number of genes. We tested the performance of the phylogenomics steps in a highly parallel versus modestly parallel architecture and found that 48-thread parallelization decreased runtime 4.2-fold on average compared to four threads (Fig. 7B). While the maximum number of species included in our test datasets (25 ingroup species) is modest in comparison to some implementations in non-plant taxa (Steenwyk et al. 2022; Tao et al. 2024), the built-in parallelization features of *ERCnet* show potential to help enable high-throughput analyses in reasonable timeframes with modest computational resources.

**Fig. 7:**
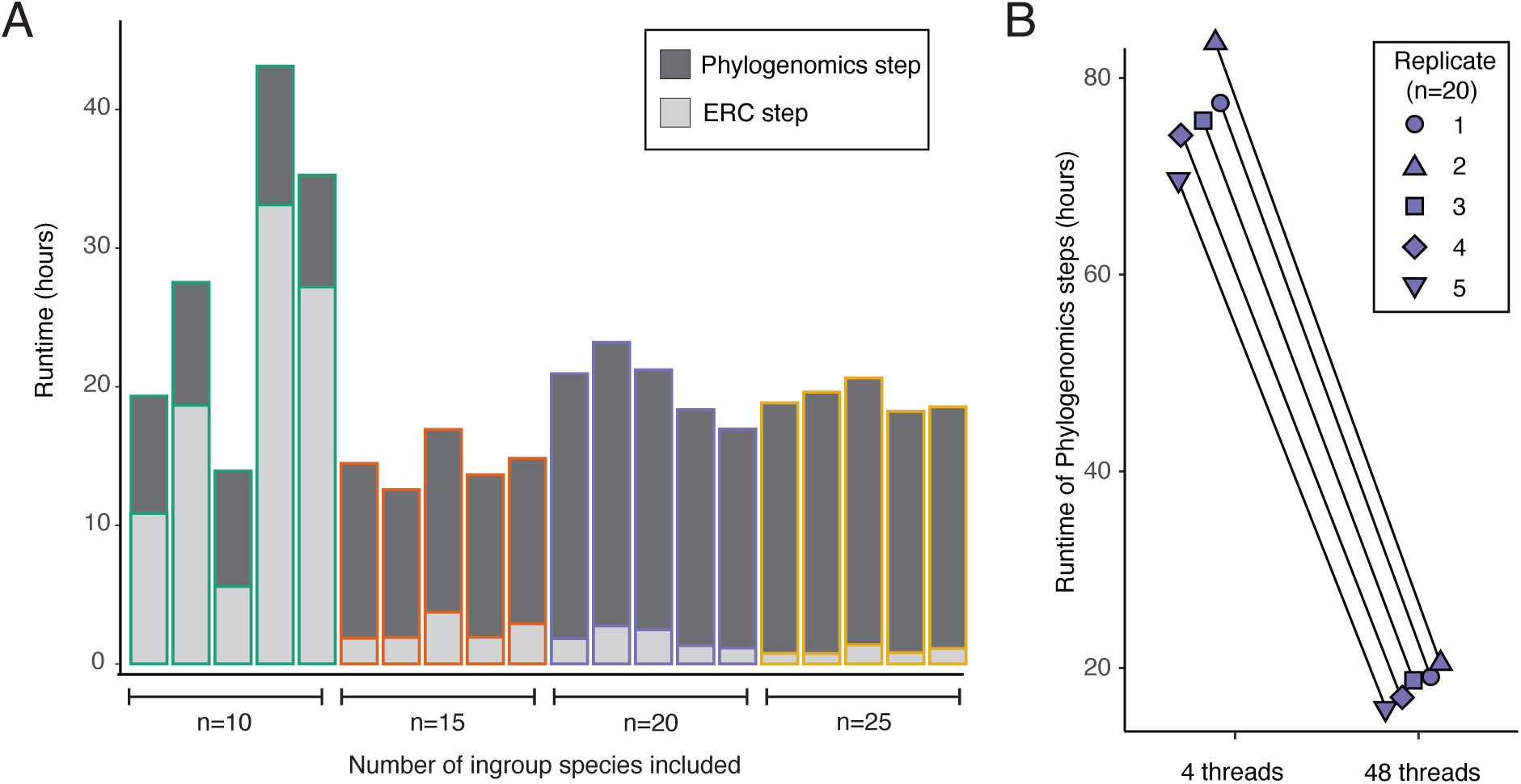
Performance of *ERCnet* using parallel processing. (A) Runtime of the analytical steps of *ERCnet* parallelized with 48 threads. The reconciliation and network analyses steps omitted because they are very fast relative to the phylogenomics steps and pairwise ERC steps and contribute negligibly to the overall runtime. (B) Runtime of the *Phylogenomics* steps of *ERCnet* on either four threads or 48 threads. The highly parallelized 48-thread runs were approximately 4-fold faster than four-thread runs.

## Discussion

### Assessing significance filters and error rates with simulations of protein evolution

Our simulations revealed *ERCnet* can achieve low rates of false positive and negative error. Using the most conservative filters, *ERCnet* achieved false positive rates below 10% in most cases for R2T and below 1% in most cases for BXB, both of which are an encouraging indication that *ERCnet* will be a useful and reliable predictive tool. However, it should be noted that our simulations were designed for parameter exploration and proof-of-principle and likely do not capture important sources of noise that impact real datasets. Developing highly realistic simulations and incorporating gold-standard plant interaction data into ERC-based networks represents an important area of future research for the field.

Our simulations revealed that our BXB method is a relatively conservative approach. This is desirable in applications in which researchers seek high-confidence interactions; however, our results demonstrate that BXB combined with the Spearman correlation method produces false negative rates approaching 100%, meaning BXB should not be combined with the Spearman method. Taken together, our results outline a useful roadmap to help users select *ERCnet* options to tailor the stringency of significance thresholds to their experimental priorities.

### Optimized design for ERC analyses

The choice of how many taxa to include is typically made in the early stages of planning an ERC study, yet the field lacks clear guidance in this area. A wide range of taxon sample sizes have been used in past studies, ranging from 18 budding yeast species (Clark et al. 2012) to 472 insect species (Tao et al. 2024). To our knowledge, past choices are mainly driven by the availability of publicly available genome/proteome sequences and are likely aimed at including the maximum possible number of taxa under the rationale that this is favorable because it increases statistical power. The datasets tested here represent only a modest range in size and extend to smaller numbers of species than have typically been used in ERC, yet we were able to detect patterns across this range that draw the ‘more is more’ rationale into question. Most notably, we demonstrate a tradeoff between the proportion of the proteome that passes filters as suitable to study via ERC and the potential statistical power of those ERC analyses gained in large datasets (Fig. 4). Our gene family filtering criteria are more lenient than the criteria typically applied in ERC studies, but even so, our largest datasets (25 ingroup species), restricted the number of gene families passing filters to <7,000 proteins (Fig. 4A), which contain only ∼30% of Arabidopsis proteins. This suggests that there is at least some benefit to limiting sampling sets to a modest number of species. Consistent with this idea, we did not observe a clear impact of dataset size on the functional clustering of ERC networks, with datasets as small as *n*=10 showing significant clustering (although with a larger degree of variance across replicate sample datasets) (Fig. 5). Indeed, branch-length measurement strategy (R2T vs BXB) may have a largest influence on *ERCnet* performance, with notable differences in error rate on simulated data (Figs. 2 and S1), network size (Fig. 5), and network overlap (Fig. 6 and Table 1). The cases in which BXB showed a substantial (Figs. 2 and S1) or incremental (Fig 5 C-D) improvement over R2T may be owed to the additional internal branch length datapoints made available in the BXB method or by the ability to reduce the statistical non-independence/pseudo-replication problem inherent to the R2T method (see Results). In any case, our results suggest that combining a moderately sized dataset with the BXB branch length method could present an optimal balance of proteome coverage and statistical power.

It should be noted that the test set of species included here were selected based on model organism status and phylogenetic representation, but not based on organismal biology or ecology features. This is a major difference from some prior studies (Williams et al. 2019; Forsythe et al. 2021) that have incorporated *a priori* knowledge of important transitions in evolutionary rates into taxon sampling decisions. The use of ERC to explore functional modules that coevolve during specific evolutionary transitions (Gatts et al. 2024) or to explore how rate variation interacts with complex categorical (Redlich et al. 2023) or quantitative trait evolution (Kowalczyk et al. 2019) is an exciting new direction (Hu et al. 2019; Treaster et al. 2023; Yan et al. 2023). In such cases, it may make sense to prioritize sampling species that represent important evolutionary transitions or trait values. Although we have not explored this dimension of taxon sampling here, it is possible that *a priori* information about phenotypic and evolutionary rate variation could alter the dynamics of how dataset size influences ERC analyses.

### Dataset dependence and consensus signatures of ERC across angiosperm datasets

There are countless biological and technical factors that could lead to a given ERC hit not being detected in a given sampling set. The highly dimensional nature of all-by-all ERC analyses mean that differences in genome quality and organismal biology can become amplified, leading to vastly inconsistent ERC-based networks. These factors could explain why the ERC hits produced by different runs of *ERCnet* show only a modest degree of overlap (Fig 6). While the level of overlap we observed clearly exceeds a random background, the observation that most ERC hits were found in only a single run raises concerns about the reproducibility of ERC results. On the other hand, the lack of overlapping hits may suggest that ERC signatures reflect the specific biology of the taxa included in an *ERCnet* analysis. This demonstrates that applying ERC in unique sets of taxa has the potential to identify novel interactions. Our results also highlight that ERC does not return all protein-protein interactions in the cell. Instead, ERC is equipped to return the interactions that have been influenced by variable selection pressures across the chosen taxa.

Given the dataset dependence of many ERC hits, it could be especially informative to identify the shortlist of ERC hits that are detected across multiple datasets and *ERCnet* runs because the identity of the proteins involved in these interactions could reveal fundamental drivers of plant molecular evolution. We find that interactions among chloroplast-localized proteins constitute a large portion of our consensus ERC hits (Table 1), including interactions among functional categories that have been documented in previous plastid-nuclear ERC work. However, the prevalence of chloroplast-related proteins in this study is especially novel and striking because the taxa used here were not selected with chloroplast function in mind. Despite this (and perhaps because of this), our consensus ERC hits revealed several new interactions that had not been identified in prior plastid-nuclear focused ERC analyses. These hits include proteins with little functional information on TAIR, meaning understanding these interactions could lead to novel insights into chloroplast biology. Beyond the novel predictions of gene function provided by our consensus ERC hits, our results raise the question: why do chloroplast-localized proteins exhibit ERC to such an exceptional degree? The answer to this question remains elusive and is outside of the scope of this study, but perhaps the consensus approach that is made possible by *ERCnet* will lay the groundwork for addressing this type of question.

In general, our consensus ERC results resemble prior attempts to identify the ‘strongest’ set of ERC hits (Forsythe et al. 2021; Gatts et al. 2024) in that our shortlist is a ‘mixed bag’ that includes some extremely intuitive predicted interactions but also pairs of proteins that seemingly lack a clear functional connection, at least based on summary information from TAIR. For example, we predict an interaction between *SHOT1-binding ATPase 4* and *SHOT1-binding ATPase 2*, which share identical TAIR descriptions, supporting co-functionality. On the other hand, our ERC hit connecting *Fibrillin10* (located in the nucleus and chloroplast; involved in structural activity) with *Ribosomal Protein UL15M* (located in the mitochondrion) does not inherently suggest co-functionality at first glance; however, this ERC hit could represent a functional pathways that spans compartments, such as proteostasis or retrograde signaling, which ERC may be adept at detecting (Forsythe et al. 2021; Gatts et al. 2024; Little et al. 2024). In any case, understanding which of these ‘surprising’ ERC hits constitute novel functional insights versus noise remains an ongoing challenge to interpretating ERC results; however, our approach of identifying ERC hits spanning randomly sampled datasets could present a valuable means of pinpointing the groups of proteins whose interaction and coevolution are indicative of the most widespread drivers of intermolecular co-evolution in plants.

## Materials and methods

### Proteome simulations

We used the python module, *pyvolve* (v. 0.9.0), to simulate protein evolution of 1000 gene families and combined the resulting protein sequences into proteomes for 21 species (Spielman and Wilke 2015). To simulate each gene family, first we generated a randomly branching bifurcating tree with each branch length randomly sampled from a distribution between 0.01 and 0.05. Trees generated in this way were used as guide trees to simulate protein sequences using *pyvovle*. Protein sequence lengths were randomly set to between 200 and 1000 amino acids. All proteins with simulated with the “jtt” model of evolution. We designated tree 1-100 as our ‘co-acceleration’ trees, which were used to simulate protein sequences that underwent accelerated evolution along specific branches of the tree. We simulated this acceleration by randomly selecting five branches on the tree and multiplying the branch length by ten. Protein families 1-100 were simulated using this accelerated tree as guide tree. Protein families 101-1000 were simulated with the original un-accelerated guide tree. After all protein families were simulated, the protein family sequences were deconstructed and moved into 21 proteome files, which were input into the standard *ERCnet* workflow (Fig. 2). For this dataset, a true-positive ERC hit was defined as a significant ERC hit in which both interaction partners were from protein families 1-100. The false positive rate was calculated as the number of significant ERC hits in which at least one interaction partner was from protein family 101-1000 divided by the total number of comparisons of that type. The false negative rate was defined was 1 minus the number of true-positives divided by the total number comparisons between pairs of proteins families from 1-100.

As a separate simulation-based test, we simulated a separate set of proteomes with a critical difference being that the five accelerated branches were randomly chosen separately for each of the 100 accelerated gene trees. While this dataset still includes 100 accelerated protein families, each is accelerated in a different set of branches, meaning they are not ‘co-accelerated’, thus serving as a negative control for *ERCnet* detection of correlated evolution (Fig. S1). In this dataset, any significant ERC hit was considered a ‘spurious’ ERC hit.

All simulations used in this study can be reproduced with the *Simulations.py* script. Calculations of false positive/negative rates can be reproduced with the *Sim_error_track.py* script. Both scripts are available on the *ERCnet* GitHub page.

### Obtaining and processing randomized plant proteome datasets

We created a pool of 35 plant proteome sequence dataset (Table S1), which included a mixture of crop species and model organisms, chosen for their general high-quality genomic resources. Sequences were obtained from the *Phytozome* plant comparative genomics portal (release 13) (Goodstein et al. 2012). We downloaded the ‘primary transcripts only’ version fasta files for each species to avoid splice isoforms being mistaken for paralogs. Fasta files were reformatted to standardize sequence ID formatting (e.g. “Atha AT1G06950” for an *Arabidopsis thaliana* protein sequence), using *SeqKit* Unix tools (Shen et al. 2016).

From our pool of 35 plant genomes, we randomly selected species for datasets of size *n*=10, *n*=15, *n*=20, and *n*=25, where *n* refers to the number of ingroup species included. We performed five replicates for each *n* value, each with its own randomly selected set of taxa. All datasets included *Amborella trichopoda* as an outgroup and *Arabidopsis thaliana* as a common focal species for cross-dataset comparisons. Three replicates of randomly selected taxa were performed for each *n* value, resulting in a total of 20 datasets, each of which was subjected to full *ERCnet* analyses.

*ERCnet* analyses require clustered gene families as input, which we obtained by running *Orthofinder* (v2.2.5) (Emms and Kelly 2015, 2019). We ran *Orthofinder* separately on each test dataset using the following command.

`orthofinder -f <path/to/dir/containing/proteomes/> -y -X -M msa -t <number of threads available on computing system>`

*Orthofinder* species tree outputs were manually inspected to verify that *Amborella trichopoda* was the outgroup. When this was not the case, we re-ran *Orthofinder* with a constrained species tree consistent with established species relationships (Jansen et al. 2007). *Orthofinder* “hierarchical orthogroups’ (HOGs) for each dataset were used as input for each *ERCnet* run.

### Setting up computing environment and installing ERCnet dependencies

*ERCnet* has been tested extensively on multiple MacOS and Linux-based computers. For this study, we ran all *ERCnet* analyses on a node on a HPC Linux server equipped with dual AMD EPYC 7713 processors, each featuring 64 physical cores with a base clock speed of 2.0 GHz, for a combined total of 128 physical cores. The system includes 1 TB (16x64 GB) of PC4-25600 3200 MHz DDR4 ECC RDIMM RAM, two 240 GB Micron 5400 PRO Series SATA SSDs for the operating system, and a 256 TB storage capacity. We created two *Conda* virtual environments to accomplish *ERCnet* steps that require *Python 3* and *Python 2*. Our environments utilized *Python* 3.9.7 and *Python* 2.7.18, respectively. The *Conda* environments, including *ERCnet* dependencies, can be recreated using the .yml files provided with the *ERCnet* package on *GitHub*. All *ERCnet* runs presented in this study were performed using *ERCnet* (v1.2.0) (see *ERCnet* documentation for release information).

### Phylogenomic analyses

The first steps of *ERCnet* are implemented in the *Phylongenomics.py* script, which filters gene families by user defined parameters (see -r and -p flag below). Next, multiple sequence alignments are generated using the L-INS-I algorithm in *MAFFT* (v7.515) (Katoh and Standley 2013). Next, *TAPER* (Zhang et al. 2021) is (optionally) used to identify outlier sequences and filter out noise from heterogeneity. *GBLOCKS* (*v0.91b)* (Castresana 2000) is then used to trim poorly aligned regions from alignments. Alignments that are shorter than 100 amino acids (set with the ‘-l 100’ flag) following trimming are discarded. Below is an example command used to call Phylogenomics.py, including descriptions of all required arguments. For detailed descriptions of optional arguments, see the *ERCnet* GitHub page.

`./Phylogenomics.py -j <Job Name> -o <OrthoFinder Results Files> -m <Threads> -s -p 4 -r 11 -T <Julia Environment Path< -n 1`

The -j flag indicates the desired name for the output folder for all *ERCnet* outputs. This ‘jobname’ is used to help *ERCnet* find needed input/output files throughout the *ERCnet* workflow. The -o flag indicates the path to *Orthofinder* results used in the analyses. The -m flag allows for parallel multithreading to speed up analysis time by using *Joblib* (see below). The -s flag allows the user to provide a custom species mapping file to help *ERCnet* assign gene names to species. The -p and -r flags are used during an initial filtering step to remove gene families with too many paralogs or too little species representation, which are unlikely to be useful in *ERCnet* analyses (e.g. gene families consisting of only 100 paralogs from one species). The -p flag indicates the maximum number of paralogs per species allowed in each gene family. For example, “-p 3” would cause *ERCnet* to exclude a gene family that contains four or more paralogs for any species. Note that *Orthofinder’s* HOGs (Emms and Kelly 2019) split deep paralogs into separate gene families, so the paralogs detected during this filtering step represent only relatively recent gene duplicates. We set -p to 3 for all *ERCnet* runs used in this study. The -r flag controls the minimum number of species represented in each gene family. For example, “-r 6” indicates that at least six unique ingroup species must be present in a gene family in order for *ERCnet* to retain the gene family. For the randomized datasets of varying size used in this study, we used set -r to the ceiling of (*n*/2)+1. For example, for n=20 dataset, r was set to 11. The -T flag is (optionally) used to tell *ERCnet* where to find the *Julia* installation used to run *TAPER.* The -n flag allows the user to specify the species tree node from which to obtain *Orthofinder* HOG subtrees (Emms and Kelly 2019).

To achieve parallelization for the steps that represent potential computational bottlenecks (Fig. 1), we use *Joblib*, an open-source *Python* library that uses *C* and *loky* to circumvent *Python’s* Global Interpreter Lock and spawn parallel worker processes.

### Gene-tree/species-tree reconciliation

Once gene trees are generated for all retained gene families, *ERCnet* performs a GT/ST reconciliation step via the *GTST_reconciliation.py* script. GTST_reconciliation.py employs *DLCpar* (v2.0.1 Kellis) to infer gene duplication, coalescence, and loss, which allows *ERCnet* to reconcile conflicting gene trees and species and perform BLR (See Results and Fig 1B). While *ERCnet* generally runs inside a *Python 3* environment, *DLCpar* requires *Python 2*. We use *Conda* environments to easily toggle between *Python 2* and *Python 3*. Once a *Python* 2 environment is activated, *GTST_reconciliation.py* can be called with the following command:

`./GTST_reconciliation.py -j <Job Name>`

The -j flag requires the same job name from the previous step to continue the analysis.

### ERC analyses with branch length reconciliation

After GT/ST reconciliation, *ERCnet* performs ERC analyses in an all-vs-all fashion by comparing the branch lengths for every possible pairwise combination of gene trees. To account for lineage-specific differences in rate of evolution, each branch length on each gene tree is normalized by the genome-wide average for the corresponding branch before correlation analyses are performed. Below is an example script used to call ERC_analysis.py.

`./ERC_analyses.py -j <Job Name> -m <Threads> -s <Focal Species> -b <Branch Reconciliation Method>`

The -j flag continues to use the same job name from previous steps. The -m flag determines how many threads used for parallelizing the analysis. The -s flag indicates which species should be the focal species for the analysis. The -b flag controls which branch length correlation method (R2T or BXB) is used. For this study, we ran *ERCnet* with both methods in order to compare results.

### Network analyses

The final step of *ERCnet* filters ERC results to retain ERC hits according to user-defined cutoffs and filtering criteria. *Network_analysis.py* includes options to filter results by R-squared values and p-values obtained from Pearson and/or Spearman correlation analyses. Users choose to filter by raw p-value or p-values that are multiple-test corrected with the Benjamini-Hochberg false discovery rate (FDR) method (Benjamini and Hochberg 1995). The script also includes options for building network graphs to visualize interaction networks. Nodes indicate gene families and edges between nodes indicate significant ERC signal. Many options for network visualization, clustering, and export are available (see *ERCnet* GitHub page). Additionally, *Network_analysis.py* provides the option to evaluate clustering of user-defined functional gene annotations by calculating the assortativity coefficient (Newman 2003) using the *igraph* package (Csardi and Nepusz 2006). *Network_analyses.py* was run with the following command:

`./Network_analyses.py -j <Job Name> -m <Branch method used> -y <Clustering algorithm> -s <Focal Species> -f <ERC results filename>`

The -j flag continues the same job name from previous steps. The -m flag informs the script which branch length method was used from the previous step. The -f flag indicates which ERC results file should be analyzed in the event that prior step was run multiple times. The -y flag allows the user to specify which clustering method they would like for generating network plots. The -s flag specifics which focal species the user is providing. The -f flag provides the script with the ERC results tsv file the user wishes to use for this portion of the analysis. Multiple ERC results files can be generated by *ERC_analysis.py*.

### Analysis of ERC hit overlap

To ask if ERC hits from multiple *ERCnet* runs overlap more than would be expected by chance, we performed a permutation test in which we randomized the ERC results table. The ERC results table includes a “Gene A” and “Gene B” corresponding to the two gene families being compared for ERC signature in a given pairwise combination. First, to gauge overall overlap between different *<u>ERCnet</u>* runs, we simply counted the number of *ERCnet* runs in which a given Gene A and Gene B combination are present among the ERC hits. To achieve a ‘null distribution’, we randomized the Gene B column of the tables and again calculated ERC hit overlap across *ERCnet* runs. We replicated this procedure ten times to get an average degree of overlap for *ERCnet* runs with different parameters. To analyze consensus ERC hits, we extracted the ERC hits present in the top three levels of overlap for R2T and BXB *ERCnet* runs (Table 1) and obtained functional annotation data for the *Arabidopsis thaliana* paralog in each gene family from TAIR (Lamesch et al. 2012).

---

**Table S1:** Proteome data used for random sample datasets.

| Species | File name | Common name | Version | Clade |
| --- | --- | --- | --- | --- |
| *Arabidopsis thaliana* | Atha | Thale cress | Araport11 | Eudicot |
| *Amborella trichopoda* | Atri | NA | v1.0 | Outgroup |
| *Ananas comosus* | Acom | Pineapple | v3 | Monocots |
| *Asparagus officinalis* | Aoff | Asparagus | V1.1 | Monocots |
| *Brachypodium distachyon* | Bdis | Brachypodium | v3.1 | Monocots |
| *Dioscorea alata* | Dala | Purple yam | v2.1 | Monocots |
| *Hordeum vulgare* | Hvul | Barley | r1 | Monocots |
| *Musa acuminata* | Macu | Banana | v1 | Monocots |
| *Miscanthus sinensis* | Msin | Chinese silver grass | v7.1 | Monocots |
| *Oryza sativa* | Osat | Rice | v7.0 | Monocots |
| *Oropetium thomaeum* | Otho | Dessication tolerant grass | v1.0 | Monocots |
| *Panicum hallii* | Phal | Hall's panic grass | v2.0 | Monocots |
| *Panicum virgatum* | Pvir | Switchgrass | v5.1 | Monocots |
| *Sorghum bicolor* | Sbic | Sorghum | v3.1.1 | Monocots |
| *Spirodela polyrhiza* | Spol | Duckweed | v2 | Monocots |
| *Setaria viridis* | Svir | Wild foxtail millet | v2.1 | Monocots |
| *Triticum aestivum* | Taes | Triticum | v2.2 | Monocots |
| *Zostera marina* | Zmar | Eelgrass | v2.2 | Monocots |
| *Zea mays* | Zmay | Maize | RefGen v4 | Monocots |
| *Arabidopsis lyrata* | Alyr | Sand cress | v2.1 | Eudicot |
| *Beta vulgaris* | Bvul | Sugar beet | EL10 1.0 | Eudicot |
| *Carica papaya* | Cpap | Papaya | ASGPBv0.4 | Eudicot |
| *Capsella rubella* | Crub | Pink shepard's purse | v1.1 | Eudicot |
| *Eutrema salsugineum* | Esal | Saltwater cress | v1.0 | Eudicot |
| *Populus trichocarpa* | Ptri | California poplar | v4.1 | Eudicot |
| *Vitis vinifera* | Vvin | Common grape vine | v2.1 | Eudicot |
| *Fragaria vesca* | Fves | Strawberry | v4.0.a2 | Eudicot |
| *Eucalyptus grandis* | Egra | Ecalyptus | v2.0 | Eudicot |
| *Helianthus annuus* | Hann | Sunflower | r1.2 | Eudicot |
| *Mimulus guttatus* | Mgut | Monkeyflower | v2.0 | Eudicot |
| *Prunus persica* | Pper | Peach | v2.1 | Eudicot |
| *Solanum lycopersicum* | Slyc | Tomato | TAG2.4 | Eudicot |
| *Spinacia oleracea* | Sole | Spinach | Spov3 | Eudicot |
| *Vigna unguiculata* | Vung | Black-eyed pea | v1.2 | Eudicot |
| *Lactuca sativa* | Lsat | Lettuce | V8 | Eudicot |

**Table S2:** Gene families filtered from sample datasets.

| Spec-<br>ies in<br>data-<br>set<br>(n) | Rep-<br>licate | Repres<br>ent-<br>ation<br>filter<br>cutoff<br>(r) | Para-<br>log<br>filter<br>cutoff<br>(p) | Number of gene families |  |  |  |  | Proportion of gene families |  |  |  |
| --- | --- | --- | --- | --- | --- | --- | --- | --- | --- | --- | --- | --- |
|  |  |  |  | Re-<br>tained | Dis-<br>qualified<br>by r | Dis-<br>qualified<br>by p | Dis-<br>qualified<br>by p and<br>r | Ortho-<br>finder<br>output | Re-<br>tained | Dis-<br>qualified<br>by r | Dis-<br>qualified<br>by p | Dis-<br>qualified<br>by p and<br>r |
| 10 | 1 | 6 | 4 | 9106 | 20191 | 2886 | 4472 | 37337 | 0.24 | 0.54 | 0.08 | 0.12 |
| 10 | 2 | 6 | 4 | 9842 | 18711 | 2249 | 4491 | 36026 | 0.27 | 0.52 | 0.06 | 0.12 |
| 10 | 3 | 6 | 4 | 8837 | 20403 | 2855 | 4251 | 36942 | 0.24 | 0.55 | 0.08 | 0.12 |
| 10 | 4 | 6 | 4 | 10982 | 20112 | 3298 | 4132 | 39108 | 0.28 | 0.51 | 0.08 | 0.11 |
| 10 | 5 | 6 | 4 | 10262 | 10161 | 1616 | 2645 | 25458 | 0.40 | 0.40 | 0.06 | 0.10 |
| 15 | 1 | 9 | 4 | 8565 | 24280 | 3238 | 4740 | 41522 | 0.21 | 0.58 | 0.08 | 0.11 |
| 15 | 2 | 9 | 4 | 7399 | 28322 | 4502 | 5516 | 46259 | 0.16 | 0.61 | 0.10 | 0.12 |
| 15 | 3 | 9 | 4 | 9391 | 22459 | 2592 | 4935 | 40124 | 0.23 | 0.56 | 0.06 | 0.12 |
| 15 | 4 | 9 | 4 | 8719 | 23148 | 3089 | 5190 | 40858 | 0.21 | 0.57 | 0.08 | 0.13 |
| 15 | 5 | 9 | 4 | 9154 | 23719 | 2586 | 4889 | 41088 | 0.22 | 0.58 | 0.06 | 0.12 |
| 20 | 1 | 11 | 4 | 8446 | 30865 | 3639 | 5982 | 49696 | 0.17 | 0.62 | 0.07 | 0.12 |
| 20 | 2 | 11 | 4 | 9422 | 21912 | 2487 | 4954 | 39622 | 0.24 | 0.55 | 0.06 | 0.13 |
| 20 | 3 | 11 | 4 | 8524 | 26066 | 3369 | 5379 | 44091 | 0.19 | 0.59 | 0.08 | 0.12 |
| 20 | 4 | 11 | 4 | 7984 | 30249 | 3686 | 6317 | 48894 | 0.16 | 0.62 | 0.08 | 0.13 |
| 20 | 5 | 11 | 4 | 7413 | 29484 | 4678 | 6257 | 48514 | 0.15 | 0.61 | 0.10 | 0.13 |
| 25 | 1 | 14 | 4 | 6571 | 32019 | 4823 | 7513 | 51627 | 0.13 | 0.62 | 0.09 | 0.15 |
| 25 | 2 | 14 | 4 | 6316 | 34568 | 5324 | 8036 | 54869 | 0.12 | 0.63 | 0.10 | 0.15 |
| 25 | 3 | 14 | 4 | 6802 | 33350 | 5114 | 7119 | 53090 | 0.13 | 0.63 | 0.10 | 0.13 |
| 25 | 4 | 14 | 4 | 6439 | 35191 | 5002 | 7704 | 54956 | 0.12 | 0.64 | 0.09 | 0.14 |
| 25 | 5 | 14 | 4 | 6622 | 31182 | 4888 | 7115 | 50521 | 0.13 | 0.62 | 0.10 | 0.14 |

**Fig. S1:**
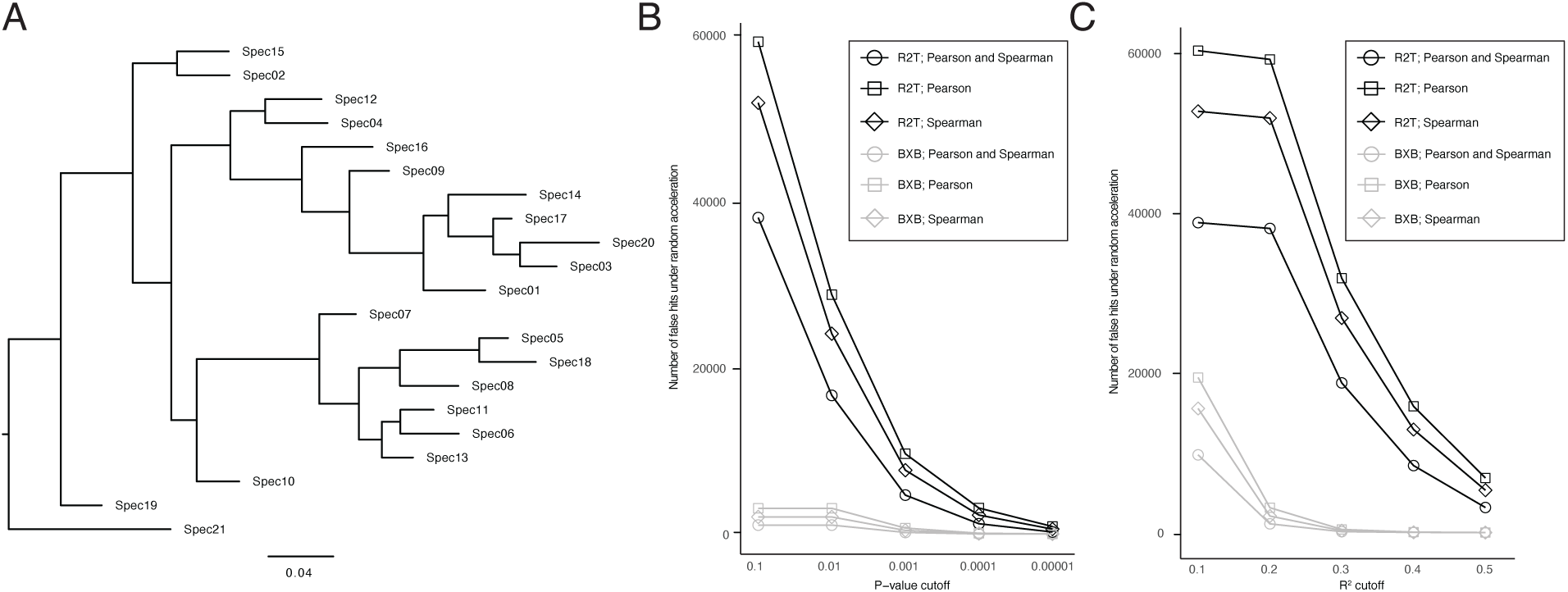
Simulations of spurious ERC hits. Simulations performed under an uncorrelated acceleration model. (A) Unaccelerated tree used as the background species tree. For each protein family simulation, five branches randomly chosen branches were accelerated, leading to uncorrelated patterns of acceleration between different protein families. (B-C) Number of ‘spurious’ ERC hits in networks obtained across different P-value (B) and *R^2^*value significance filtering cutoffs.

**Fig. S2:**
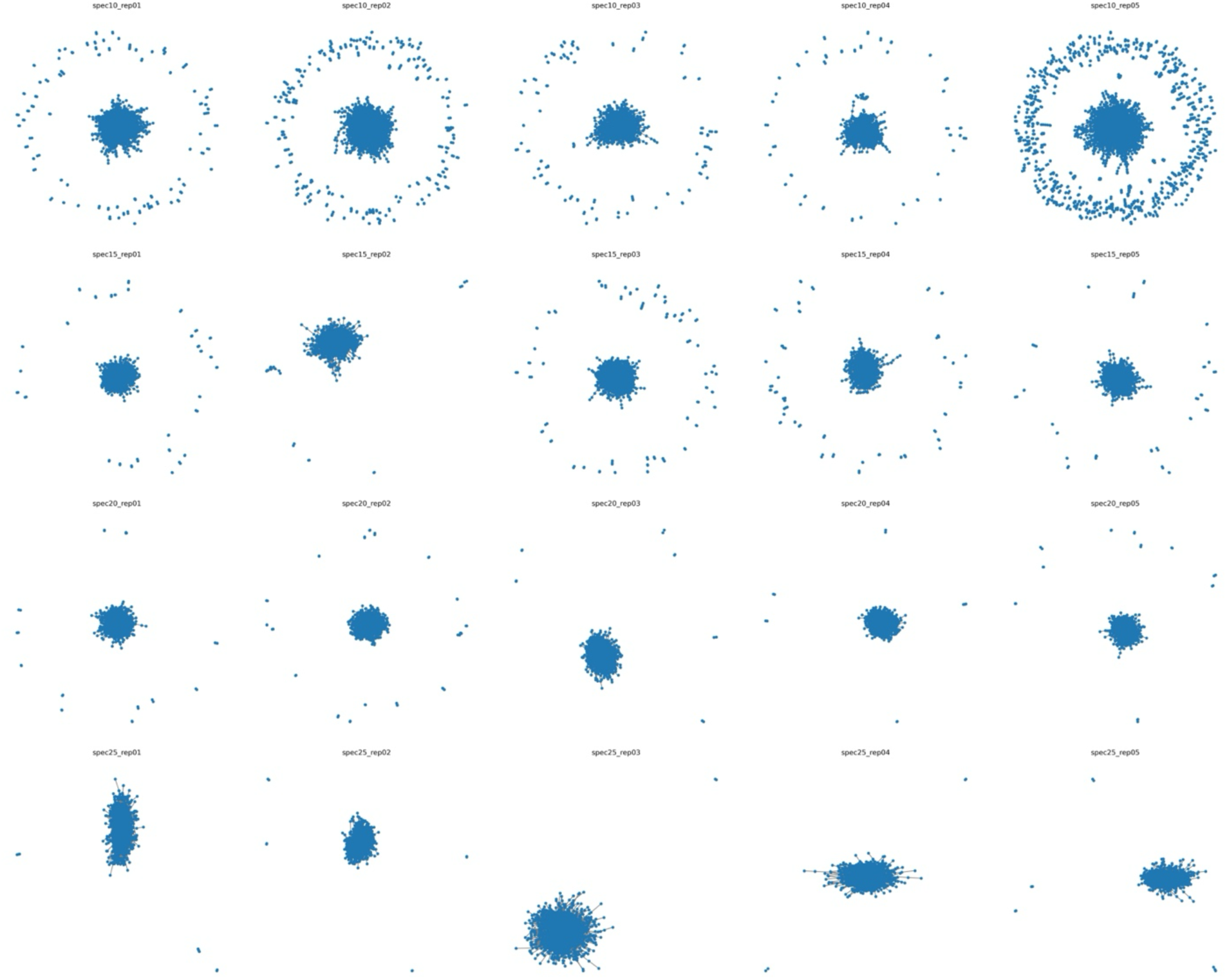
Network graphs for R2T-based networks. Network graphs from empirical replicate datasets obtained using the root-to-tip method. Number of ingroup species and replicate number are indicated by “spec” and “rep”, respectively.

**Fig. S3:**
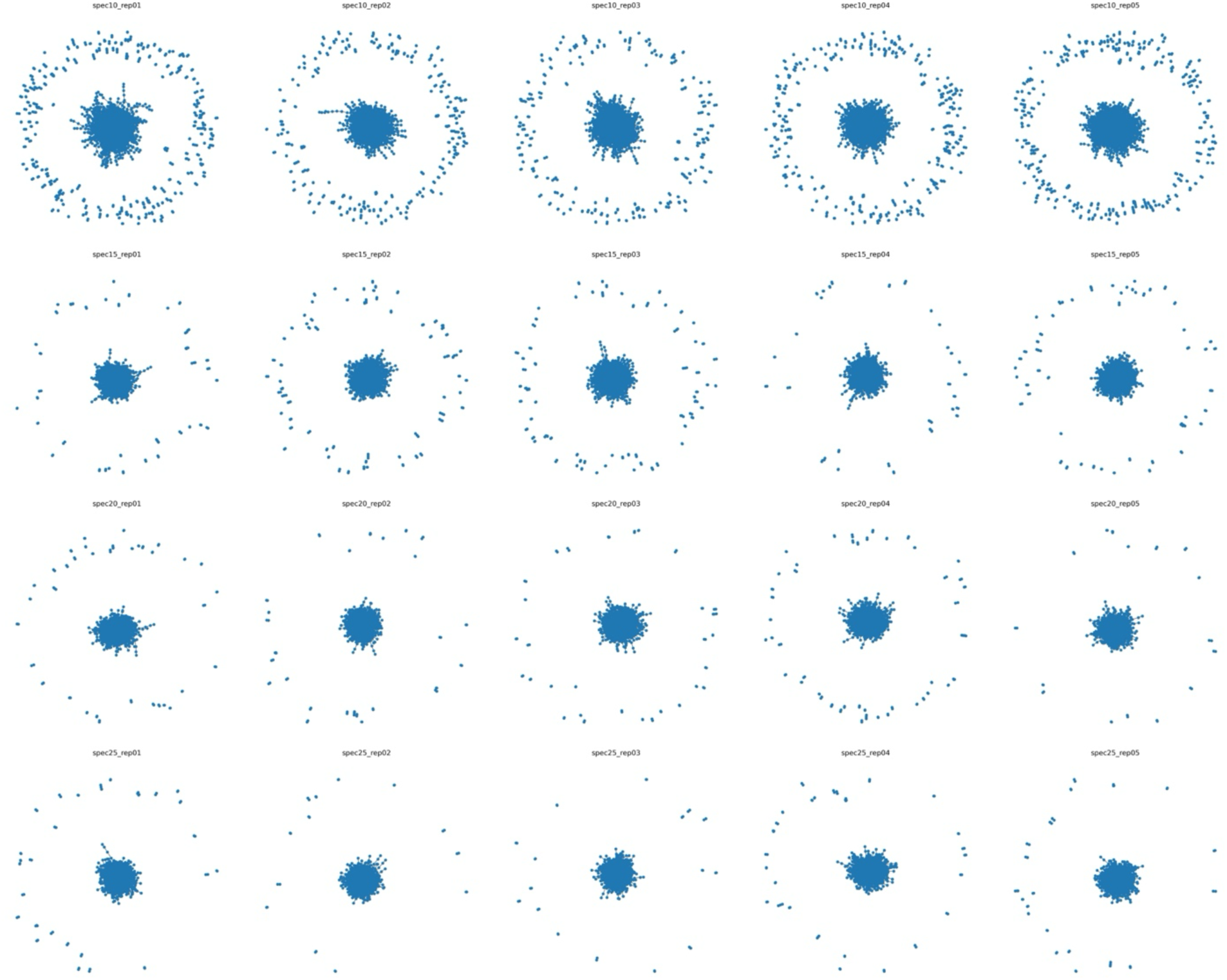
Network graphs for BXB-based networks. Network graphs from empirical replicate datasets obtained using the branch-by-branch method. Number of ingroup species and replicate number are indicated by “spec” and “rep”, respectively.

